# Speech-induced suppression and vocal feedback sensitivity in human cortex

**DOI:** 10.1101/2023.12.08.570736

**Authors:** Muge Ozker, Leyao Yu, Patricia Dugan, Werner Doyle, Daniel Friedman, Orrin Devinsky, Adeen Flinker

**Author notes:** **Corresponding Author:** Muge Ozker, **e-mail:**.

## Abstract

Across the animal kingdom, neural responses in the auditory cortex are suppressed during vocalization, and humans are no exception. A common hypothesis is that suppression increases sensitivity to auditory feedback, enabling the detection of vocalization errors. This hypothesis has been previously confirmed in non-human primates, however a direct link between auditory suppression and sensitivity in human speech monitoring remains elusive. To address this issue, we obtained intracranial electroencephalography (iEEG) recordings from 35 neurosurgical participants during speech production. We first characterized the detailed topography of auditory suppression, which varied across superior temporal gyrus (STG). Next, we performed a delayed auditory feedback (DAF) task to determine whether the suppressed sites were also sensitive to auditory feedback alterations. Indeed, overlapping sites showed enhanced responses to feedback, indicating sensitivity. Importantly, there was a strong correlation between the degree of auditory suppression and feedback sensitivity, suggesting suppression might be a key mechanism that underlies speech monitoring. Further, we found that when participants produced speech with simultaneous auditory feedback, posterior STG was selectively activated if participants were engaged in a DAF paradigm, suggesting that increased attentional load can modulate auditory feedback sensitivity.

## Introduction

A major question in neuroscience is how do animals distinguish between stimuli originating from the environment and those produced by their own actions. Sensorimotor circuits share a common mechanism across the animal kingdom in which sensory responses to self-generated motor actions are suppressed. It is commonly hypothesized that suppressing responses to predicted self-generated stimuli increases sensitivity of the sensory system to external stimuli. (Poulet and Hedwig 2002, Poulet and Hedwig 2006, Crapse and Sommer 2008, E.Vonholstan, Glenn et al. 2011, Schneider and Mooney 2018). Furthermore, it enables detection and correction of motor errors by providing a template of the predicted sensory outcome to compare with the actual sensory outcome. In the domain of speech, this mechanism is described in models which suggest that neural responses in the auditory cortex are suppressed during speech production. When there is a mismatch between the predicted auditory outcome and the actual auditory feedback, responses in the auditory regions are enhanced to encode the mismatch and inform vocal-motor regions to correct vocalization (Hickok, Houde et al. 2011, Houde and Nagarajan 2011, Tourville and Guenther 2011).

A common experimental strategy to generate mismatch between the predicted auditory outcome and the actual auditory feedback is to perturb auditory feedback during speech production. Auditory feedback perturbations are usually applied either by delaying auditory feedback (DAF), which disrupts speech fluency (Lee 1950, Fairbanks 1955, Stuart, Kalinowski et al. 2002), or by shifting voice pitch and formants, which result in compensatory vocal changes in the opposite direction of the shift (Houde and Jordan 1998, Jones and Munhall 2000, Niziolek and Guenther 2013). Numerous electrophysiological and neuroimaging studies investigated neural responses during speech production both in the absence and presence of auditory feedback perturbations. In support of speech production models, these studies have repeatedly reported suppressed responses in auditory cortex during speaking compared with passive listening to speech (Numminen, Salmelin et al. 1999, Wise, Greene et al. 1999, Curio, Neuloh et al. 2000, Houde, Nagarajan et al. 2002, Christoffels, Formisano et al. 2007, Ford, Roach et al. 2010, Niziolek, Nagarajan et al. 2013), as well as enhanced responses when auditory feedback was perturbed indicating sensitivity to auditory feedback (Tourville, Reilly et al. 2008, Behroozmand, Karvelis et al. 2009, Chang, Niziolek et al. 2013, Greenlee, Behroozmand et al. 2013, Kort, Nagarajan et al. 2014, Behroozmand, Shebek et al. 2015, Ozker, Doyle et al. 2022). However, it is not clear whether the same or distinct neural populations in the auditory cortex show speech-induced suppression and sensitivity to auditory feedback.

While auditory responses are largely suppressed during speech production, detailed investigations using neurosurgical recordings revealed that the degree of suppression was variable across cortical sites, and auditory cortex also exhibited non-suppressed and enhanced responses (albeit less common) (Creutzfeldt and Ojemann 1989, Flinker, Chang et al. 2010, Greenlee, Jackson et al. 2011), mirroring results from non-human primate studies using single unit recordings (Eliades and Wang 2003, Eliades and Wang 2008). In the same non-human primate study, it was reported that neurons that were suppressed during vocalization showed increased activity when auditory feedback was perturbed (Eliades and Wang 2008). Based on this finding, we predicted that if speech-induced suppression enables detection and correction of speech errors, suppressed auditory sites should be sensitive to auditory feedback, thus exhibit enhanced neural responses to feedback perturbations. Alternatively, if suppression and speech monitoring are unrelated processes, then suppressed sites should be distinct from the ones that are sensitive to auditory feedback.

The level of attention during speech monitoring can vary depending on the speech task. During normal speech production, speech monitoring does not require a conscious effort, however it is a controlled, attentional process during an auditory feedback perturbation task (Hashimoto and Sakai 2003). It is well known that selective attention enhances auditory responses and improves speech perception under noisy listening conditions or when multiple speech streams are present (Mesgarani and Chang 2012, Golumbic, Ding et al. 2013). We predicted that increased attention to auditory feedback under adverse speaking conditions, such as during an auditory feedback perturbation task, should increase feedback sensitivity and elicit larger responses in the auditory cortex compared to normal speech production.

To summarize, in this study we aimed to test the hypothesis that speech-induced suppression increases sensitivity to auditory feedback in human neurophysiological recordings. We predicted that auditory sites showing speech induced suppression would elicit enhanced responses to auditory feedback perturbations. Further, we aimed to investigate the role of attention in auditory feedback sensitivity by comparing auditory responses during an auditory feedback perturbation task compared with normal speech production.

To address these aims, we used iEEG recordings in neurosurgical participants, which offers a level of spatial detail and temporal precision that would not be possible to achieve using non-invasive techniques. We first identified the sites that show auditory suppression during speech production, and then employed a DAF paradigm to test whether the same sites show sensitivity to perturbed feedback. Our results revealed that overlapping sites in the STG exhibited both speech-induced auditory suppression and sensitivity to auditory feedback with a strong correlation between the two measures, supporting the hypothesis that auditory suppression predicts sensitivity to speech errors in humans. Further, we showed that auditory responses in the posterior STG are enhanced in a DAF task compared to normal speech production, even for trials in which participants receive simultaneous auditory feedback (no-delay condition). This result suggests that increased attention during an auditory feedback perturbation task can modulate auditory feedback sensitivity and posterior STG is a critical region for this attentional modulation.

## Materials and Methods

### Participant Information

All experimental procedures were approved by the New York University School of Medicine Institutional Review Board. 35 neurosurgical epilepsy patients (19 females, mean age: 31, 23 left, 9 right and 3 bilateral hemisphere coverage) implanted with subdural and depth electrodes provided informed consent to participate in the research protocol. Electrode implantation and location were guided solely by clinical requirements. 3 patients were consented separately for higher density clinical grid implantation, which provided denser sampling of underlying cortex.

### Intracranial Electroencephalography (iEEG) Recording

iEEG was recorded from implanted subdural platinum-iridium electrodes embedded in flexible silicon sheets (2.3 mm diameter exposed surface, 8 x 8 grid arrays and 4 to 12 contact linear strips, 10 mm center-to-center spacing, Ad-Tech Medical Instrument, Racine, WI) and penetrating depth electrodes (1.1 mm diameter, 5-10 mm center-to-center spacing 1 x 8 or 1 x 12 contacts, Ad-Tech Medical Instrument, Racine, WI). 3 participants consented to a research hybrid grid implanted which included 64 additional electrodes between the standard clinical contacts (16 × 8 grid with sixty-four 2 mm macro contacts at 8 x 8 orientation and sixty-four 1 mm micro contacts in between, providing 10 mm center-to-center spacing between macro contacts and 5 mm center-to-center spacing between micro/macro contacts, PMT corporation, Chanassen, MN). Recordings were made using one of two amplifier types: NicoletOne amplifier (Natus Neurologics, Middleton, WI), bandpass filtered from 0.16-250 Hz and digitized at 512 Hz. Neuroworks Quantum Amplifier (Natus Biomedical, Appleton, WI) recorded at 2048 Hz, bandpass filtered at 0.01 to 682.67 Hz and then downsampled to 512 Hz. A two-contact subdural strip facing toward the skull near the craniotomy site was used as a reference for recording and a similar two-contact strip screwed to the skull was used for the instrument ground. iEEG and experimental signals (trigger pulses that mark the appearance of visual stimuli on the screen, microphone signal from speech recordings and feedback voice signal) were acquired simultaneously by the EEG amplifier in order to provide a fully synchronized dataset.

## Experimental Design

### Experiment 1: Auditory word repetition (AWR)

35 participants performed the experiment. Stimuli consisted of 50 items (nouns) taken from the revised Snodgrass and Vanderwart object pictorial set (e.g. “drum’, “hat”, “pencil”) (Rossion and Pourtois 2004, Shum, Fanda et al. 2020). Auditory words presented randomly (2 repetitions) through speakers. Participants were instructed to listen to the presented words and repeat them out loud at each trial.

### Experiment 2: Visual word reading (VWR)

The same 35 participants performed the experiment. Stimuli consisted of the same 50 words used in Experiment 1, however visually presented as text stimuli on the screen in a random order (2 repetitions). Participants were instructed to read the presented word out loud at each trial.

### Experiment 3: Delayed auditory feedback (DAF)

A subgroup of 14 participants performed this experiment. Stimuli consisted of 10 different 3-syllable words visually presented as text stimuli on the screen (e.g. “envelope”, “umbrella”, “violin”). Participants were instructed to read the presented word out loud at each trial. As participants spoke, their voices were recorded using the laptop’s internal microphone, delayed at 4 different amounts (no-delay, 50, 100, 200ms) using custom script (MATLAB, Psychtoolbox-3) and played back to them through earphones. Trials, which consisted of different stimulus-delay combinations, were presented randomly (3 to 8 repetitions). Behavioral and neural data from the DAF experiment were used in a previous publication from our group (Ozker, Doyle et al. 2022).

### Experiment 4: Visual word reading with auditory feedback (VWR-AF)

A subgroup of 4 participants performed an additional visual word reading experiment, in which they were presented with the word stimuli as in Experiment 3 and heard their simultaneous (no-delay) voice feedback through earphones.

### Statistical Analysis

Electrodes were examined for speech related activity defined as significant high gamma broadband responses. Unpaired t-tests were performed to compare responses to a baseline for each electrode and multiple comparisons were corrected using the false discovery rate (FDR) method (q=0.05). Electrodes that showed significant response increase (p < 10^-4^) either before (−0.5 to 0 s) or after speech onset (0 to 0.5 s) with respect to a baseline period (−1 to −0.6 s) and at the same time had a large signal-to-noise ratio (μ/σ > 0.7) during either of these time windows were selected. Electrode selection was first performed for each task separately, then electrodes that were commonly selected were further analyzed. For the analysis of the DAF experiment, one-way ANOVA was calculated using the average neural response as a dependent variable and feedback delay as a factor to assess the statistical significance of response enhancement in a single electrode.

### Experiment Setup

Participants were tested while resting in their hospital bed in the epilepsy-monitoring unit. Visual stimuli were presented on a laptop screen positioned at a comfortable distance from the participant. Auditory stimuli were presented through speakers in the AWR and VWR experiments and through earphones (Bed Phones On-Ear Sleep Headphones Generation 3) in the DAF and in the VWR-AF experiment. Participants were instructed to speak at a normal voice level and sidetone volume was adjusted to a comfortable level at the beginning of the DAF experiment. DAF and VWR-AF experiments were performed consecutively and sidetone volume was kept the same in the two experiments. Participants’ voice was recorded using an external microphone (Zoom H1 Handy Recorder). A TTL pulse marking the onset of a stimulus, the microphone signal (what the participant spoke) and the feedback voice signal (what the participant heard) were fed in to the EEG amplifier as an auxiliary input in order to acquire them in sync with EEG samples. Sound files recorded by the external microphone were used for voice intensity analysis. Average voice intensity for each trial was calculated in dB using the ‘Intensity’ object in Praat software (Boersma 2001).

### Electrode Localization

Electrode localization in individual space as well as MNI space was based on co-registering a preoperative (no electrodes) and postoperative (with electrodes) structural MRI (in some cases a postoperative CT was employed depending on clinical requirements) using a rigid-body transformation. Electrodes were then projected to the surface of cortex (preoperative segmented surface) to correct for edema induced shifts following previous procedures (Yang, Wang et al. 2012) (registration to MNI space was based on a non-linear DARTEL algorithm (Ashburner 2007). Within participant anatomical locations of electrodes was based on the automated FreeSurfer segmentation of the participant’s pre-operative MRI. We recorded from a total of 3591 subdural and 1361 depth electrode contacts in 35 participants. Subdural electrode coverage extended over lateral temporal, frontal, parietal and lateral occipital cortices. Depth electrodes covered additional regions to a limited extent including the transverse temporal gyrus, insula and fusiform gyrus. Contacts that were localized to the cortical white matter were excluded from the analysis. To categorize electrodes in the STG into anterior and posterior groups, lateral termination of the transverse temporal sulcus was used as an anatomical landmark (Greenlee, Jackson et al. 2011, Nourski, Steinschneider et al. 2016).

### Neural Data Analysis

Electrodes with epileptiform activity or artifacts caused by line noise, poor contact with cortex and high amplitude shifts were removed from further analysis. A common average reference was calculated by subtracting the average signal across all electrodes from each individual electrode’s signal (after rejection of electrodes with artifacts). The analysis of the electrophysiologic signals focused on changes in broadband high gamma activity (70–150 Hz). To quantify changes in the high gamma range, the data were bandpass filtered between 70 and 150 Hz, and then a Hilbert transform was applied to obtain the analytic amplitude.

Recordings from the DAF and VWR-AF experiments were analyzed using the multitaper technique, which yields a more sensitive estimate of the power spectrum with lower variance, thus is more beneficial when comparing neural responses to incremental changes in stimuli. Continuous data streams from each channel were epoched into trials (from −1.5 s to 3.5 s with respect to speech onset). Line noise at 60, 120 and 180 Hz were filtered out. 3 Slepian tapers were applied in timesteps of 10 ms and frequency steps of 5 Hz, using temporal smoothing (tw) of 200 ms and frequency smoothing (fw) of ±10 Hz. Tapered signals were then transformed to time-frequency space using discrete Fourier transform and power estimates from different tapers were combined (MATLAB, FieldTrip toolbox). The number of tapers (K) were determined by the Shannon number according to the formula: K=2*tw*fw-1 (Percival and Walden 1993). The high gamma broadband response (70-150 Hz) at each time point following stimulus onset was measured as the percent signal change from baseline, with the baseline calculated over all trials in a time window from −500 to −100 ms before stimulus onset.

### Suppression Index (SuppI) Calculation

Suppression of neural activity is measured by comparing responses in two time periods in the AWR task. First time period was during listening the stimulus (0-0.5 s) and the second time period was during speaking (0-0.5 s). For each trial, average responses over Listen and Speak periods were found and suppression was measured by calculating Listen-Speak/Listen+Speak. Then suppression values were averaged across trials to calculate a single suppression index for each electrode. For the neural activity, raw high gamma broadband signal power was used instead of the percent signal change to ensure that the suppression index values varied between −1 to 1, indicating a range from complete enhancement to complete suppression respectively.

### Sensitivity Index (SensI) Calculation

Sensitivity to DAF is measured by comparing neural responses to increasing amounts of feedback delay. Neural responses in each trial were averaged in a time period following the voice feedback (0-0.5 s). For each electrode, a sensitivity index was calculated by measuring the trial-by-trial Spearman correlation between the delay condition and the averaged neural response. A large sensitivity value indicated a strong response enhancement with increasing delays.

## Results

In order to assess cortical responses during perception and production of speech, and quantify speech-induced auditory suppression, participants (N = 35) performed an auditory word repetition (AWR) task. We examined the response patterns in seven different cortical regions including superior temporal gyrus (STG), middle temporal gyrus (MTG), supramarginal gyrus (SMG), inferior frontal gyrus (IFG), middle frontal gyrus (MFG), precentral gyrus (preCG) and postcentral gyrus (postCG) (**Fig 1A**). As an index of the neural response, we used the high gamma broadband signal (70-150 Hz, *see Methods*), which correlates with the spiking activity of the underlying neuronal population (Mukamel, Gelbard et al. 2005, Crone, Sinai et al. 2006, Cardin, Carlen et al. 2009, Ray and Maunsell 2011, Lachaux, Axmacher et al. 2012).

**Figure 1.**
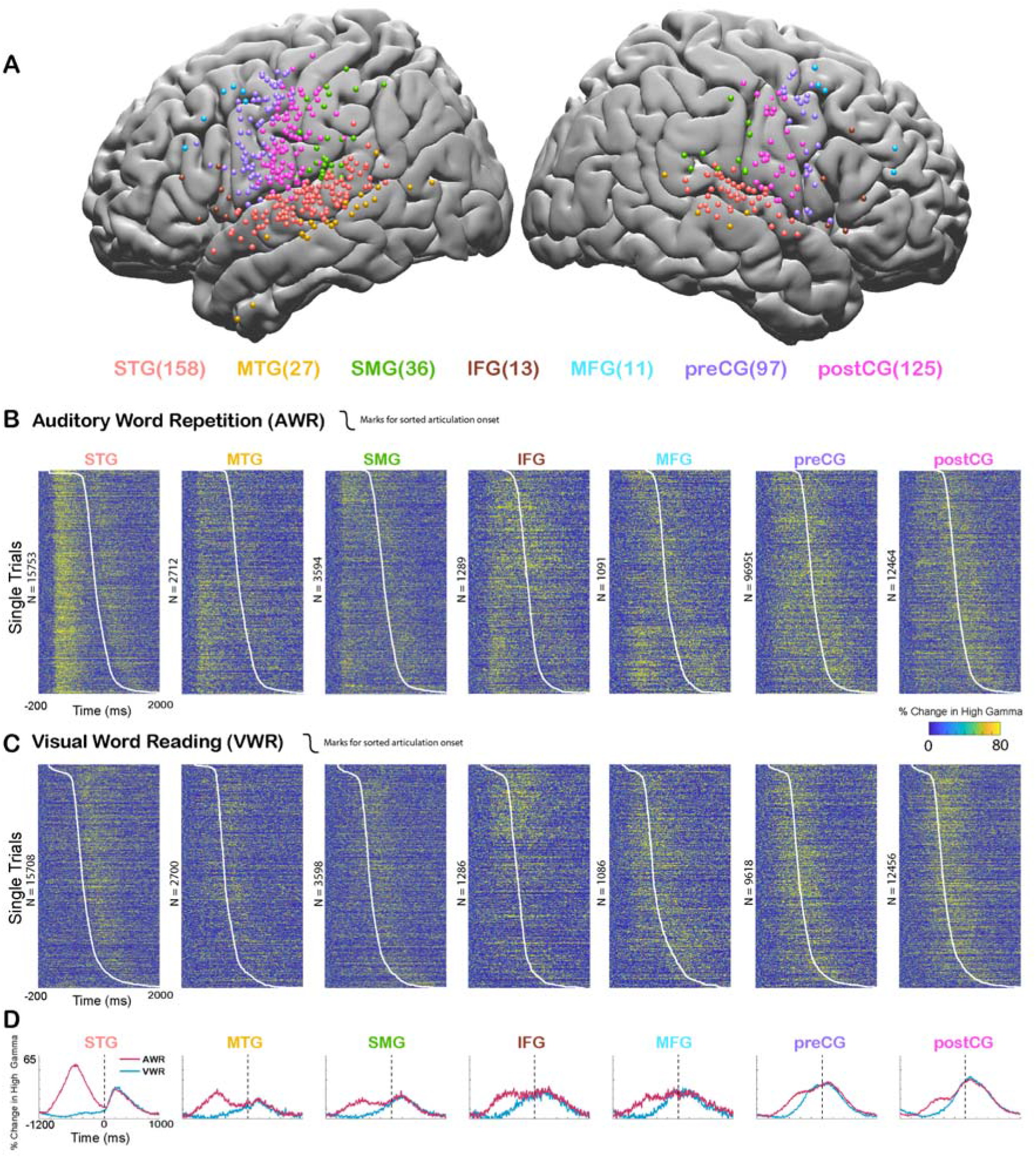
Cortical responses during speech tasks. **A.** Electrodes from all participants (n = 35) are shown on a template brain with different colors corresponding to different regions (number of electrodes in each region denoted in the parentheses). **B.** High gamma broadband responses (70-150 Hz) for individual trials in an Auditory Word Repetition task are shown for each region. **C.** High gamma responses for individual trials in a Visual Word Reading task are shown for each region. Trials are sorted with respect to speech onset (white line). **D.** Mean high gamma broadband response averaged across trials are shown for each region with the width representing the standard error of the mean across electrodes. Time=0 indicates speech production onset.

We analyzed the responses in two different time windows: During passive listening of the auditory stimulus (0-500 ms after stimulus onset) and during speaking when participants repeated the perceived auditory stimulus (0-500 ms after articulation onset). Average responses were larger during passive listening in STG (Average % signal change ± SEM; Listen: 62.1±0.6, Speak: 29.8±0.4), MTG (32.7±0.9, 22.3±0.9) and SMG (27.4±0.8, 25.8±0.7) compared with speaking. Conversely, responses were larger during speaking in IFG (29.2±1.3, 31.2±1.3), MFG (28.3±1.6, 31.4±1.3), preCG (27.4±0.4, 37±0.5) and postCG (26±0.4, 42±0.5). These results suggested that auditory regions responded more strongly during passive listening compared to speaking, verifying previous reports of neural response suppression to self-generated speech in auditory cortex (**Fig 1B-D**).

In the AWR task, participants heard the same auditory stimulus twice in each trial, once from a recorded female voice and once from their own voice. It is well known that repeated presentation of a stimulus results in the suppression of neural activity in regions that process that stimulus, a neural adaptation phenomenon referred to as repetition suppression (Grill-Spector, Henson et al. 2006, Todorovic and de Lange 2012). To ensure that our observed suppression of neural activity in auditory regions was not due to repetition suppression, but rather was induced by speech production, we performed a visual word reading (VWR) task, in which participants hear the auditory stimulus only once (from their own voice). Response magnitudes during speaking in the AWR and VWR tasks were similar (paired t-test: t (466) = 0.62, p = 0.53), characterized by a strong correlation across electrodes (Pearson’s Correlation: r = 0.9006, p = 0). These results suggested that repetition of the auditory stimulus in the AWR task did not affect response magnitudes and the observed reduction in response magnitudes was induced by speech production.

To quantify the amount of speech-induced suppression, we calculated a Suppression Index (SuppI) for each electrode by comparing neural responses during listening versus speaking in the AWR task (SuppI = Listen-Speak/Listen+Speak; *see Methods*). A positive SuppI indicated a response suppression during speaking compared to listening and was observed most strongly in middle to posterior parts of STG, followed by MTG and SMG. A negative SuppI indicated a response enhancement during speaking compared to listening and was observed in motor regions, most strongly in the postCG (**Fig 2A-B**).

**Figure 2.**
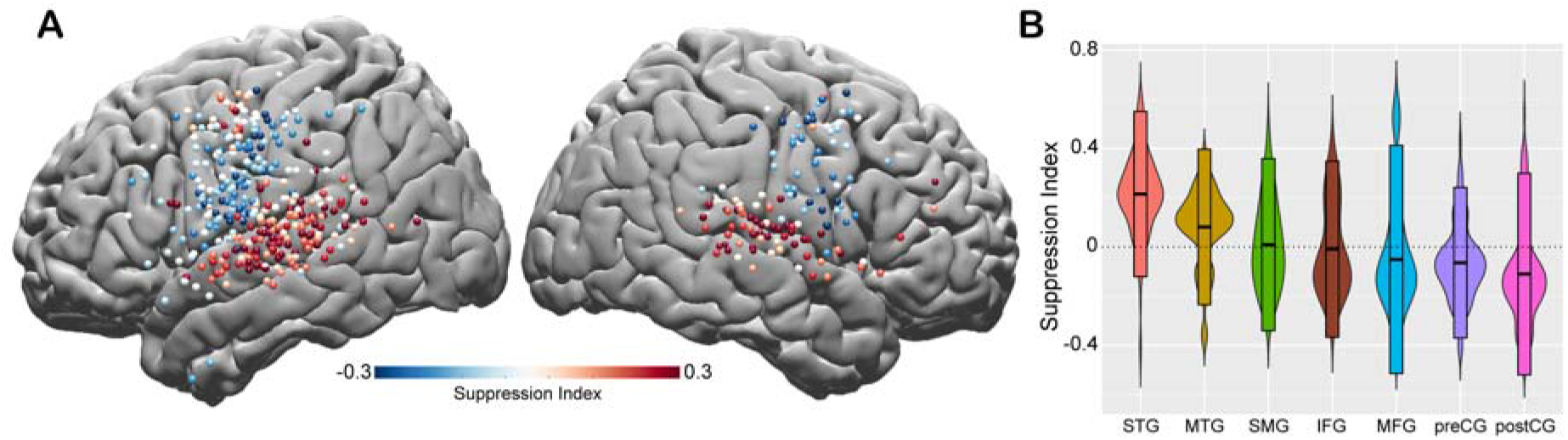
Spatial topography of speech-induced auditory suppression. **A.** Suppression indices for all electrodes are shown on a template brain. Red color tones indicate smaller neural activity during speaking, while blue electrodes indicate larger neural activity during speaking compared to listening in the Auditory Word Repetition task. **B.** Suppression indices averaged across electrodes are shown for each region sorted from largest to smallest mean suppression index. Boxplots indicate mean ± SD.

After mapping the topographical distribution of suppression indices across the cortex, we focused on understanding the functional role of auditory suppression in speech monitoring. We hypothesized that the degree of speech-induced auditory suppression should be tightly linked to sensitivity to speech errors, as predicted by current models (Houde and Nagarajan 2011, Tourville and Guenther 2011) and neural data in non-human primates (Eliades and Wang 2008). To test this hypothesis, we used an additional task, in which we delayed the auditory feedback (DAF) during speech production to disrupt speech fluency. In this task, 14 participants repeated the VWR task while they were presented with their voice feedback through earphones either simultaneously (no-delay) or with a delay (50, 100 and 200 ms; *see Methods*). In a previous study (Ozker, Doyle et al. 2022), using the same data set, we demonstrated that participants slowed down their speech in response to DAF (Articulation duration; DAF_0_: 0.698, DAF_50_: 0.726, DAF_100:_ 0.737, and DAF_200:_ 0.749 milliseconds). Moreover, auditory regions exhibited an enhanced response that varied as a function of feedback delay, likely representing an auditory error signal encoding the mismatch between the expected and the actual feedback. However, those results were not directly linked to auditory suppression.

Here, we compared neural responses in the AWR and the DAF tasks to test whether auditory regions that exhibit strong speech-induced suppression also exhibit large auditory error responses to DAF, which would indicate strong sensitivity to speech errors. In a single participant, we demonstrated that a representative electrode on the STG with strong auditory suppression (Average % signal change in 0-500 ms; Listen: 124±7, Speak: 20±3, SuppI: 0.27) exhibited significant response enhancement (DAF_0_: 135±12, DAF_50_: 134±8, DAF_100_: 175±10, DAF_200_: 208±17, ANOVA: F (3, 116) = 8.5, p = 3.7e-05) (**Fig 3A-B**), while a nearby electrode with weaker auditory suppression (Listen: 116±6, Speak: 80±4, SuppI: 0.06) did not exhibit significant response enhancement with feedback delays (DAF_0_: 360±29, DAF_50_: 328±24, DAF_100_: 379±31, DAF_200_: 419±30, ANOVA: F (3, 116) = 1.73, p = 0.16) (**Fig 3C-D**).

**Figure 3.**
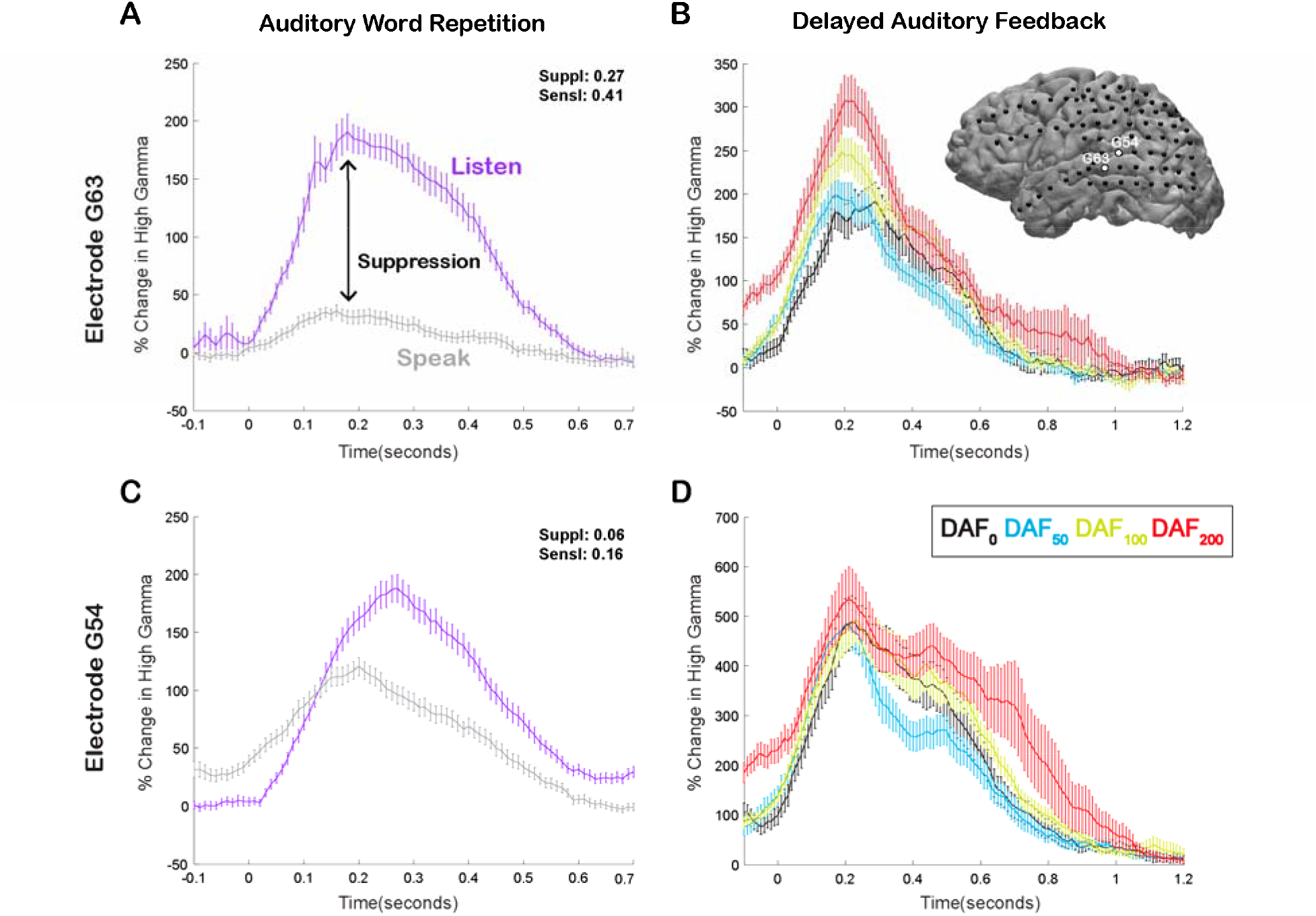
Speech-induced auditory suppression and sensitivity to delayed auditory feedback in representative electrodes in a single participant. **A.** High gamma broadband response (70-150 Hz) in electrode G63 showing a large amount of auditory suppression during speaking words compared to listening to the same words. Error bars indicate SEM over trials. **B.** High gamma responses in electrode G63 to articulation of words with DAF. 0 seconds indicate the onset of the perceived auditory feedback. Inset figure shows the cortical surface model of the left hemisphere brain of a single participant. Black circles indicate the implanted electrodes. White highlighted electrodes are located on the middle (G63) and caudal (G54) STG. **C.** High gamma response in electrode G54 showing a small degree of auditory suppression during speaking words compared to listening. **D.** High gamma response in electrode G54 locked to articulation of words during DAF. 0 seconds indicate the onset of the perceived auditory feedback.

To quantify the auditory error response and measure the sensitivity of a cortical region to DAF, we calculated a Sensitivity Index (SensI) for each electrode by correlating the delay condition and the average neural response across trials (*see Methods*). A large SensI indicated a strong response enhancement (large auditory error response) with increasing delays. The degree of both speech-induced suppression and sensitivity to DAF were highly variable across the cortex, SuppI ranging from −0.46 to 0.53 and SensI ranging from −0.62 to 0.70. The largest suppression and sensitivity indices as well as a strong overlap between the two measures were observed in the STG, suggesting that auditory electrodes that show speech-induced suppression are also sensitive to auditory feedback perturbations (**Fig 4A-C**). We validated this relationship by revealing a significant correlation between suppression and sensitivity indices of auditory electrodes (n = 57, Pearson’s Correlation: r = 0.4006, p = 0.002) supporting our hypothesis and providing evidence for a common neural mechanism (**Fig 4D**).

**Figure 4.**
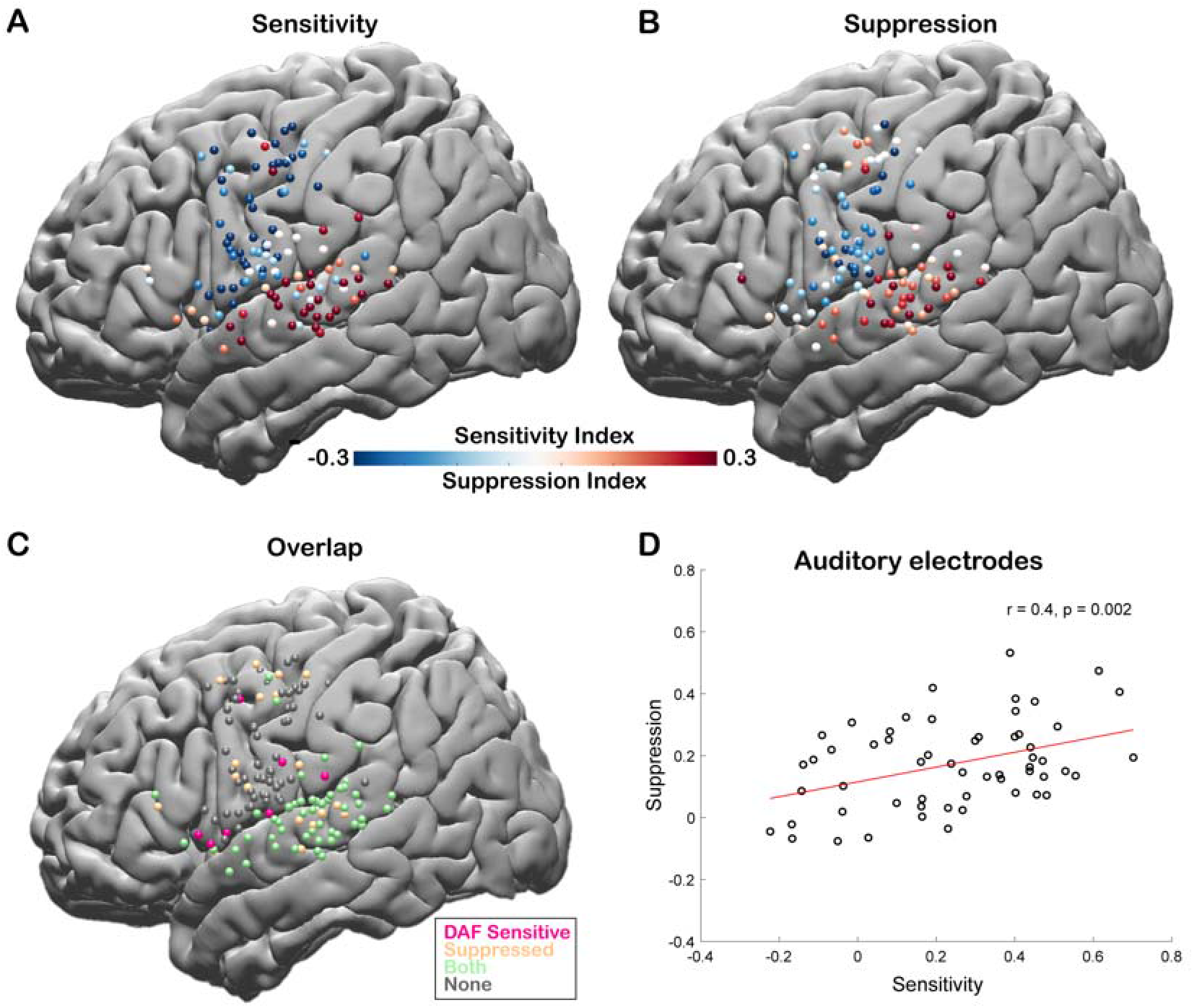
Correlation between speech-induced auditory suppression and sensitivity to delayed auditory feedback. **A.** Sensitivity indices for all electrodes are shown on a template brain (both right and left hemisphere electrodes were shown on the left hemisphere). Red tones indicate larger neural activity to increasing amount of delays in the Delayed Auditory Feedback task, while blue tones indicate the opposite. **B.** Suppression indices for all electrodes are shown on a template brain. Red tones indicate larger neural activity during listening compared to speaking in the Auditory Word Repetition task, while blue tones indicate the opposite. **C.** Electrodes that show either sensitivity to delayed auditory feedback (positive SensI value) or speech-induced auditory suppression (positive SuppI value), or both are shown on a template brain. **D.** Scatter plot and fitted regression showing a significant correlation between sensitivity to DAF and speech-induced auditory suppression across auditory electrodes. Each circle represents an electrode’s sensitivity and suppression index.

Our neural analysis revealed that response magnitudes in auditory cortex were much larger when participants heard their simultaneous voice feedback in a DAF paradigm compared with producing speech without any feedback (DAF_0_: no-delay trials) (Average % signal change in 0-500 ms; DAF_0_: 113±14, VWR: 41±7, *compare gray lines in* ***Fig 3A*** *and **C** with black lines in **Fig3B** and **D,** respectively*). We were interested in dissociating if these larger responses were merely an effect of perceiving voice feedback through earphones instead of air or rather were specific to our DAF design, likely due to increased attentional demands. Therefore, 4 participants performed an additional visual word reading task in which they were presented with their simultaneous voice feedback through earphones (VWR-AF). As previous studies have reported that DAF can increase voice intensity (Yates 1963, Howell and Archer 1984), we first verified whether participants spoke louder during the DAF task. A comparison of their voice intensity between DAF_0_ (no-delay trials in the DAF task) and the VWR-AF (standard word reading with simultaneous feedback through earphones) conditions did not show a significant difference (Voice intensity; DAF_0_: 50±11 dB, VWR: 49±12 dB; paired t-test: t (118) = 1.8, p = 0.08). After verifying that the sound volume entering the auditory system is not statistically different in the two conditions, we compared the responses in the auditory cortex and found that overall response magnitudes were now on par across the two conditions (DAF_0_: 89±17, VWR-AF: 82±17, **Fig 5A**). However, a detailed inspection of individual electrode responses revealed that some electrodes showed larger response to DAF_0_, while others showed either larger responses to VWR-AF or similar responses to both conditions (**Fig 5B**). In a single participant, we demonstrated that adjacent electrodes in the STG that are only 5 mm apart exhibited completely different response patterns. Electrodes in the more posterior parts of STG showed larger responses to DAF_0_, while electrodes in more anterior parts showed similar responses to DAF_0_ and VWR-AF (**Fig 5C**). To determine an anatomical landmark at which the reversal of response patterns occurred in the STG, we used the lateral termination of the transverse temporal sulcus (TTS) (Greenlee, Jackson et al. 2011, Nourski, Steinschneider et al. 2016) based on the individual FreeSurfer segmentation of the participant’s pre-operative MRI. Across participants, this landmark corresponded to y coordinate = −22±2.

**Figure 5.**
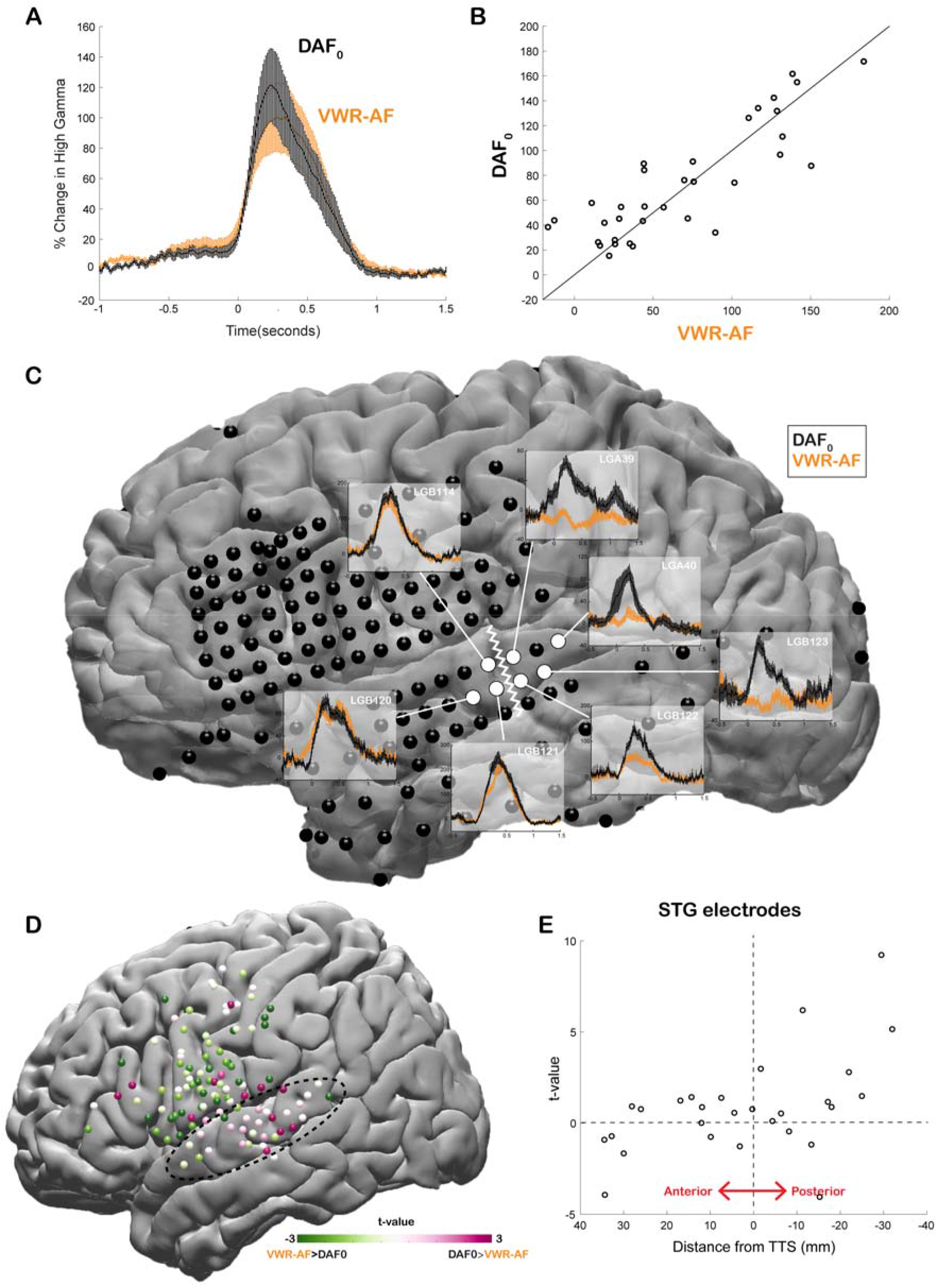
Effect of the delayed auditory feedback paradigm on neural responses during speech. **A.** High gamma broadband responses (70-150 Hz) averaged across auditory electrodes are similar during no-delay condition in the delayed auditory feedback task (DAF_0_) and during visual word reading with auditory feedback (VWR-AF). Error bars indicate SEM across electrodes. **B.** Scatter plot shows averaged high gamma responses (0-500 ms) for VWR-AF versus DAF_0_ conditions for auditory electrodes. **C.** High gamma responses for DAF_0_ and VWR-AF are shown in representative auditory electrodes in a single participant. Electrodes that are posteriorly located on the STG show larger responses to DAF_0_ condition, while electrodes that are anteriorly located on the STG show similar responses to the two conditions. The lateral termination of the transverse temporal sulcus (TTS) is identified as a landmark (white zigzagged line) that separates the two different response patterns. **D.** High gamma responses for DAF_0_ and VWR conditions were compared and resulting t-values are shown for all electrodes on a template brain. Pink color tones indicate larger responses to DAF_0_, while green color tones indicate larger responses to VWR condition. **E.** T-values calculated by comparing responses to DAF_0_ and VWR conditions are shown for all auditory electrodes with respect to their anterior-to-posterior positions to the TTS.

Next, we compared the response patterns in the two conditions for all electrodes across participants by calculating a t-value for each electrode (unpaired t-test: average responses from −200 to 500 ms). We demonstrated that auditory regions in posterior STG showed larger responses to DAF_0_ condition, while frontal motor regions showed larger responses to VWR-AF (**Fig 5D**). Lastly, we examined STG electrodes alone, sorted by their anterior-to-posterior positions with respect to the TTS. In line with the results from the single participant, electrodes that were located posteriorly within a 1 cm distance from this anatomical landmark showed significantly larger responses to the DAF_0_ condition (**Fig 5E**). These results suggest that posterior STG is more activated when participants are engaged in a speech production task that requires increased effort and attention.

## Discussion

Our study provides a detailed topographical investigation of speech-induced auditory suppression in a large cohort of neurosurgical participants. We found that while the strongest auditory suppression was observed in the STG, the degree of suppression was highly variable across different recording sites. To explain this variability, we considered the functional role of auditory suppression in speech monitoring. We showed that delaying auditory feedback during speech production enhanced auditory responses in the STG. The degree of sensitivity to feedback delays was also variable across different recording sites. We found a significant correlation between speech-induced suppression and feedback sensitivity, providing evidence for a shared mechanism between auditory suppression and speech monitoring. While there was no anatomical organization for auditory suppression and feedback sensitivity in the STG, we found an anterior-posterior organization for the effect of attention on feedback sensitivity. Auditory sites that lie posterior to the lateral termination of the TTS in the STG showed stronger activation during the DAF task compared to a standard word reading task, even for trials in which participants received simultaneous feedback, demonstrating attentional modulation of feedback sensitivity.

We observed the strongest speech-induced suppression in the middle and posterior parts of the STG. In line with previous iEEG studies, we found that degree of suppression was variable across different recording sites in the STG without any anatomical organization (Flinker, Chang et al. 2010, Greenlee, Jackson et al. 2011, Nourski, Steinschneider et al. 2016). So far, a clear gradient for speech-induced suppression has never been reported in the STG but only in the Heschl’s gyrus (HG) and superior temporal sulcus (STS) by studies that used comprehensive depth electrode coverage within the temporal lobe (Nourski, Steinschneider et al. 2016, Nourski, Steinschneider et al. 2021).

We found only a few sites with speech-induced enhancement and several sites with no response change. Based on single unit recordings in non-human primates, it is known that majority of neurons in the non-core auditory cortex exhibits suppression, while a smaller group exhibits excitation during vocalization. It is difficult to isolate speech-induced enhancement in human studies, because measurements reflect the average response of the underlying neural population, which is dominated by suppressed responses. A previous non-human primate study suggested that there might be a division of labor between the suppressed and excited neurons. They showed that when an external auditory stimulus is presented concurrently during vocalization, neurons that showed vocalization-induced suppression did not respond to the external stimulus. In contrary, neurons that showed vocalization-induced excitation responded even more when external stimulus is concurrently presented during vocalization, suggesting a role in maintaining sensitivity to the external acoustic environment (Eliades and Wang 2003). In humans there might be a similar division of labor between auditory sites that were suppressed and non-suppressed, such that while suppressed sites are engaged in monitoring self-generated sounds, non-suppressed sites maintain sensitivity to external sounds. But unfortunately, our study did not include the necessary experimental conditions to directly test this hypothesis.

Our broad topographical search using subdural electrodes revealed additional sites outside the canonical auditory regions in the STG that showed speech-induced suppression, mainly in the MTG, and a few others in the SMG and preCG. Sensorimotor regions in the preCG including inferior frontal and premotor cortices are known to activate during passive listening tasks (Wilson, Saygin et al. 2004, Pulvermuller, Huss et al. 2006, Cogan, Thesen et al. 2014), and show tuning to different acoustic properties of speech similar to the auditory regions in the STG (Mesgarani, Cheung et al. 2014, Cheung, Hamiton et al. 2016). Our results showed that isolated sites in these frontal motor regions were sensitive to DAF, confirming their auditory properties and suggesting their involvement in speech monitoring.

Current models of speech motor control predicted a shared mechanism between auditory suppression and sensitivity to speech errors suggesting a role for auditory suppression in speech monitoring (Houde and Nagarajan 2011, Tourville and Guenther 2011). Behavioral evidence in human studies showed that when auditory feedback is delayed in real time, speakers attempt to reset or slow down their speech (Lee 1950, Fairbanks 1955, Stuart, Kalinowski et al. 2002). Similarly, when fundamental frequency (pitch) or formant frequencies of the voice are shifted, speakers change their vocal output in the opposite direction of the shift to compensate for the spectral perturbation (Houde and Jordan 1998, Jones and Munhall 2000, Niziolek and Guenther 2013). Neurosurgical recordings and neuroimaging studies that investigate the brain mechanism of auditory feedback processing demonstrated that these feedback-induced vocal adjustments are accompanied by enhanced neural responses in various auditory regions (Tourville, Reilly et al. 2008, Behroozmand, Karvelis et al. 2009, Behroozmand, Shebek et al. 2015, Ozker, Doyle et al. 2022). However, it has not been clear whether it is the same or different neural populations that exhibit speech-induced suppression and enhanced responses to auditory feedback perturbations. Only in a non-human primate study, which recorded single-unit activity in auditory neurons of marmoset monkeys, it was shown that neurons that were suppressed during vocalization exhibited increased activity during frequency-shifted feedback (Eliades and Wang 2008). In contrast, to replicate this finding in humans, a previous iEEG study by Chang et al. (Chang, Niziolek et al. 2013) used frequency-shifted feedback during vowel production and found that most suppressed auditory sites did not overlap with those sensitive to feedback alterations. Using DAF instead of frequency-shifted feedback, we demonstrated a significant overlap of two neural populations in the STG, along with a strong correlation between the degree of speech-induced suppression and sensitivity to auditory feedback. This discrepancy may be due to different methods of calculating sensitivity to altered feedback. In our study, sensitivity was determined by comparing responses to delayed and non-delayed feedback during production, whereas Chang et al. compared perturbed feedback responses during production and listening. One possibility is that our approach identifies a larger auditory neural population in the STG sensitive to altered feedback. Alternatively, it could indicate a larger population highly sensitive to temporal rather than spectral perturbations in auditory feedback. Thus, we observe a wide overlap of the two neural populations in the STG showing both speech-induced suppression and sensitivity to auditory feedback. Replaying a recording of the participants’ own delayed voice back to them, which we were unable to complete in this study, would have made the results of the two studies more comparable while also completely eliminating the possibility of a sensory explanation for the observed response enhancement.

Forward models of speech production suggest that a mismatch between the predicted and the actual auditory feedback is encoded by a response enhancement in the auditory cortex signifying an error signal (Houde and Nagarajan 2011, Tourville and Guenther 2011, Hickok 2012). Our results suggested that attention to one’s own speech stream during adverse speaking conditions, such as during an auditory feedback perturbations task, might also contribute to the response enhancement in the auditory cortex. Auditory feedback control of speech was thought to be involuntary and not subject to attentional control, because several previous studies showed that participants produced compensatory responses to pitch shifts even when they were told to ignore feedback perturbations (Munhall, MacDonald et al. 2009, Zarate, Wood et al. 2010, Keough, Hawco et al. 2013). However, prolonging pitch shift duration resulted in an early vocal response that opposes the pitch shift direction and a later vocal response that follows the pitch shift direction suggesting an interplay between reflexive and top-down processes in controlling voice pitch (Hain, Burnett et al. 2000, Burnett and Larson 2002). More recent EEG studies demonstrated that dividing attention between auditory feedback and additional visual stimuli or increasing the attentional load of the task affected vocal responses as well as the magnitude of ERP components, suggesting that attention modulates auditory feedback control on both a behavioral and a cortical level (Tumber, Scheerer et al. 2014, Hu, Liu et al. 2015, Liu, Hu et al. 2015, Liu, Fan et al. 2018). In our study, we found that neural responses in the posterior STG were larger for DAF_0_ (randomly presented simultaneous feedback condition in the DAF task) as compared with the VWR-AF condition (consistent simultaneous feedback throughout standard word reading task), even though participants displayed similar vocal behavior in these two conditions. In light of the previous literature, we interpret these response differences as arising from an attentional load difference between the two tasks. In the DAF experiment, the auditory feedback was not consistent since no-delay trials were randomized with delay trials. This randomized structure of the paradigm with interleaved long delay trials (causing slowed speech) required conscious effort for speech-monitoring and thus sustained attention. While remaining cautious about this interpretation and our study’s limitation in attentional controls, we believe that this response enhancement represents an increased neural gain driven by attention to auditory feedback (Hillyard, Vogel et al. 1998), and highlights the critical role of the posterior STG in auditory-motor integration during speech monitoring (Hickok and Poeppel 2000), with its close proximity to the human ventral attention network comprising temporoparietal junction (TPJ) (Vossel, Geng et al. 2014). We leave it to future studies to include additional conditions to manipulate the direction and load of attention to further validate the influence of attention on speech monitoring.

## References

Ashburner, J. (2007). “A fast diffeomorphic image registration algorithm.” Neuroimage 38(1): 95–113.

Behroozmand, R., L. Karvelis, H. Liu and C. R. Larson (2009). “Vocalization-induced enhancement of the auditory cortex responsiveness during voice F0 feedback perturbation.” Clin Neurophysiol 120(7): 1303–1312.

Behroozmand, R., R. Shebek, D. R. Hansen, H. Oya, D. A. Robin, M. A. Howard, 3rd and J. D. Greenlee (2015). “Sensory-motor networks involved in speech production and motor control: an fMRI study.” Neuroimage 109: 418–428.

Boersma, P. (2001). “Praat, a system for doing phonetics by computer..” Glot International 5:**9/10**: 341–345.

Burnett, T. A. and C. R. Larson (2002). “Early pitch-shift response is active in both steady and dynamic voice pitch control.” The Journal of the Acoustical Society of America 112(3): 1058–1063.

Cardin, J. A., M. Carlen, K. Meletis, U. Knoblich, F. Zhang, K. Deisseroth, L. H. Tsai and C. I. Moore (2009). “Driving fast-spiking cells induces gamma rhythm and controls sensory responses.” Nature 459(7247): 663–667.

Chang, E. F., C. A. Niziolek, R. T. Knight, S. S. Nagarajan and J. F. Houde (2013). “Human cortical sensorimotor network underlying feedback control of vocal pitch.” Proc Natl Acad Sci U S A 110(7): 2653–2658.

Cheung, C., L. S. Hamiton, K. Johnson and E. F. Chang (2016). “The auditory representation of speech sounds in human motor cortex.” Elife 5.

Cogan, G. B., T. Thesen, C. Carlson, W. Doyle, O. Devinsky and B. Pesaran (2014). “Sensory-motor transformations for speech occur bilaterally.” Nature 507(7490): 94–98.

Crapse, T. B. and M. A. Sommer (2008). “Corollary discharge across the animal kingdom.” Nature Reviews Neuroscience 9(8): 587–600.

Creutzfeldt, O. and G. Ojemann (1989). “Neuronal activity in the human lateral temporal lobe. III. Activity changes during music.” Exp Brain Res 77(3): 490–498.

Crone, N. E., A. Sinai and A. Korzeniewska (2006). “High-frequency gamma oscillations and human brain mapping with electrocorticography.” Prog Brain Res 159: 275–295.

E. Vonholstan, D., H. M. Glenn and Mittelstaedt (2011). The Principle of Reafference : Interactions Between the Central Nervous System and the Peripheral Organs.

Eliades, S. J. and X. Wang (2003). “Sensory-motor interaction in the primate auditory cortex during self-initiated vocalizations.” Journal of neurophysiology 89(4): 2194–2207.

Eliades, S. J. and X. Wang (2008). “Neural substrates of vocalization feedback monitoring in primate auditory cortex.” Nature 453(7198): 1102–1106.

Fairbanks, G. (1955). “Selective vocal effects of delayed auditory feedback.” Journal of speech and hearing disorders 20(4): 333–346.

Flinker, A., E. F. Chang, H. E. Kirsch, N. M. Barbaro, N. E. Crone and R. T. Knight (2010). “Single-trial speech suppression of auditory cortex activity in humans.” Journal of Neuroscience 30(49): 16643–16650.

Greenlee, J. D., A. W. Jackson, F. Chen, C. R. Larson, H. Oya, H. Kawasaki, H. Chen and M. A. Howard III (2011). “Human auditory cortical activation during self-vocalization.” PloS one 6(3): e14744.

Grill-Spector, K., R. Henson and A. Martin (2006). “Repetition and the brain: neural models of stimulus-specific effects.” Trends Cogn Sci 10(1): 14–23.

Hain, T. C., T. A. Burnett, S. Kiran, C. R. Larson, S. Singh and M. K. Kenney (2000). “Instructing subjects to make a voluntary response reveals the presence of two components to the audio-vocal reflex.” Experimental Brain Research 130(2): 133–141.

Hashimoto, Y. and K. L. Sakai (2003). “Brain activations during conscious self-monitoring of speech production with delayed auditory feedback: an fMRI study.” Hum Brain Mapp 20(1): 22–28.

Hickok, G. (2012). “Computational neuroanatomy of speech production.” Nat Rev Neurosci 13(2): 135–145.

Hickok, G., J. Houde and F. Rong (2011). “Sensorimotor integration in speech processing: computational basis and neural organization.” Neuron 69(3): 407–422.

Hickok, G. and D. Poeppel (2000). “Towards a functional neuroanatomy of speech perception.” Trends Cogn Sci 4(4): 131–138.

Hillyard, S. A., E. K. Vogel and S. J. Luck (1998). “Sensory gain control (amplification) as a mechanism of selective attention: electrophysiological and neuroimaging evidence.” Philos Trans R Soc Lond B Biol Sci 353(1373): 1257–1270.

Houde, J. F. and M. I. Jordan (1998). “Sensorimotor adaptation in speech production.” Science 279(5354): 1213–1216.

Houde, J. F. and S. S. Nagarajan (2011). “Speech production as state feedback control.” Frontiers in human neuroscience 5: 82.

Houde, J. F., S. S. Nagarajan, K. Sekihara and M. M. Merzenich (2002). “Modulation of the auditory cortex during speech: an MEG study.” J Cogn Neurosci 14(8): 1125–1138.

Hu, H., Y. Liu, Z. Guo, W. Li, P. Liu, S. Chen and H. Liu (2015). “Attention modulates cortical processing of pitch feedback errors in voice control.” Scientific Reports 5(1): 1–8.

Jones, J. A. and K. G. Munhall (2000). “Perceptual calibration of F0 production: evidence from feedback perturbation.” J Acoust Soc Am 108(3 Pt 1): 1246–1251.

Keough, D., C. Hawco and J. A. Jones (2013). “Auditory-motor adaptation to frequency-altered auditory feedback occurs when participants ignore feedback.” BMC Neuroscience 14(1): 25.

Kort, N. S., S. S. Nagarajan and J. F. Houde (2014). “A bilateral cortical network responds to pitch perturbations in speech feedback.” Neuroimage 86: 525–535.

Lachaux, J. P., N. Axmacher, F. Mormann, E. Halgren and N. E. Crone (2012). “High-frequency neural activity and human cognition: past, present and possible future of intracranial EEG research.” Prog Neurobiol 98(3): 279–301.

Lee, B. S. (1950). “Effects of delayed speech feedback.” The Journal of the Acoustical Society of America 22(6): 824–826.

Liu, Y., H. Fan, J. Li, J. A. Jones, P. Liu, B. Zhang and H. Liu (2018). “Auditory-motor control of vocal production during divided attention: behavioral and ERP correlates.” Frontiers in Neuroscience 12: 113.

Liu, Y., H. Hu, J. A. Jones, Z. Guo, W. Li, X. Chen, P. Liu and H. Liu (2015). “Selective and divided attention modulates auditory-vocal integration in the processing of pitch feedback errors.” Eur J Neurosci 42(3): 1895–1904.

Mesgarani, N., C. Cheung, K. Johnson and E. F. Chang (2014). “Phonetic feature encoding in human superior temporal gyrus.” Science 343(6174): 1006–1010.

Mukamel, R., H. Gelbard, A. Arieli, U. Hasson, I. Fried and R. Malach (2005). “Coupling between neuronal firing, field potentials, and FMRI in human auditory cortex.” Science 309(5736): 951–954.

Munhall, K. G., E. N. MacDonald, S. K. Byrne and I. Johnsrude (2009). “Talkers alter vowel production in response to real-time formant perturbation even when instructed not to compensate.” The Journal of the Acoustical Society of America 125(1): 384–390.

Niziolek, C. A. and F. H. Guenther (2013). “Vowel category boundaries enhance cortical and behavioral responses to speech feedback alterations.” J Neurosci 33(29): 12090–12098.

Nourski, K. V., M. Steinschneider and A. E. Rhone (2016). “Electrocorticographic Activation within Human Auditory Cortex during Dialog-Based Language and Cognitive Testing.” Front Hum Neurosci 10: 202.

Nourski, K. V., M. Steinschneider, A. E. Rhone, C. K. Kovach, M. I. Banks, B. M. Krause, H. Kawasaki and M. A. Howard (2021). “Electrophysiology of the Human Superior Temporal Sulcus during Speech Processing.” Cereb Cortex 31(2): 1131–1148.

Numminen, J., R. Salmelin and R. Hari (1999). “Subject’s own speech reduces reactivity of the human auditory cortex.” Neurosci Lett 265(2): 119–122.

Ozker, M., W. Doyle, O. Devinsky and A. Flinker (2022). “A cortical network processes auditory error signals during human speech production to maintain fluency.” PLoS Biol 20(2): e3001493.

Percival, D. B. and A. T. Walden (1993). Spectral analysis for physical applications, cambridge university press.

Poulet, J. F. and B. Hedwig (2002). “A corollary discharge maintains auditory sensitivity during sound production.” Nature 418(6900): 872–876.

Poulet, J. F. and B. Hedwig (2006). “The cellular basis of a corollary discharge.” Science 311(5760): 518–522.

Pulvermuller, F., M. Huss, F. Kherif, F. Moscoso del Prado Martin, O. Hauk and Y. Shtyrov (2006). “Motor cortex maps articulatory features of speech sounds.” Proc Natl Acad Sci U S A 103(20): 7865–7870.

Ray, S. and J. H. Maunsell (2011). “Different origins of gamma rhythm and high-gamma activity in macaque visual cortex.” PLoS Biol 9(4): e1000610.

Rossion, B. and G. Pourtois (2004). “Revisiting Snodgrass and Vanderwart’s Object Pictorial Set: The Role of Surface Detail in Basic-Level Object Recognition.” Perception 33(2): 217–236.

Schneider, D. M. and R. Mooney (2018). “How movement modulates hearing.” Annual review of neuroscience 41: 553–572.

Shum, J., L. Fanda, P. Dugan, W. K. Doyle, O. Devinsky and A. Flinker (2020). “Neural correlates of sign language production revealed by electrocorticography.” Neurology 95(21): e2880–e2889.

Stuart, A., J. Kalinowski, M. P. Rastatter and K. Lynch (2002). “Effect of delayed auditory feedback on normal speakers at two speech rates.” J Acoust Soc Am 111(5 Pt 1): 2237–2241.

Todorovic, A. and F. P. de Lange (2012). “Repetition suppression and expectation suppression are dissociable in time in early auditory evoked fields.” J Neurosci 32(39): 13389–13395.

Tourville, J. A. and F. H. Guenther (2011). “The DIVA model: A neural theory of speech acquisition and production.” Lang Cogn Process 26(7): 952–981.

Tourville, J. A., K. J. Reilly and F. H. Guenther (2008). “Neural mechanisms underlying auditory feedback control of speech.” Neuroimage 39(3): 1429–1443.

Tumber, A. K., N. E. Scheerer and J. A. Jones (2014). “Attentional demands influence vocal compensations to pitch errors heard in auditory feedback.” PLoS One 9(10): e109968.

Vossel, S., J. J. Geng and G. R. Fink (2014). “Dorsal and ventral attention systems: distinct neural circuits but collaborative roles.” Neuroscientist 20(2): 150–159.

Wilson, S. M., A. P. Saygin, M. I. Sereno and M. Iacoboni (2004). “Listening to speech activates motor areas involved in speech production.” Nat Neurosci 7(7): 701–702.

Wise, R. J., J. Greene, C. Buchel and S. K. Scott (1999). “Brain regions involved in articulation.” Lancet 353(9158): 1057–1061.

Yang, A. I., X. Wang, W. K. Doyle, E. Halgren, C. Carlson, T. L. Belcher, S. S. Cash, O. Devinsky and T. Thesen (2012). “Localization of dense intracranial electrode arrays using magnetic resonance imaging.” Neuroimage 63(1): 157–165.

Zarate, J. M., S. Wood and R. J. Zatorre (2010). “Neural networks involved in voluntary and involuntary vocal pitch regulation in experienced singers.” Neuropsychologia 48(2): 607–618.

Boersma, P. (2001). “Praat, a system for doing phonetics by computer..” Glot International 5:9/10: 341–345.

Christoffels, I. K., E. Formisano and N. O. Schiller (2007). “Neural correlates of verbal feedback processing: an fMRI study employing overt speech.” Hum Brain Mapp 28(9): 868–879.

Curio, G., G. Neuloh, J. Numminen, V. Jousmaki and R. Hari (2000). “Speaking modifies voice-evoked activity in the human auditory cortex.” Hum Brain Mapp 9(4): 183–191.

Ford, J. M., B. J. Roach and D. H. Mathalon (2010). “Assessing corollary discharge in humans using noninvasive neurophysiological methods.” Nat Protoc 5(6): 1160–1168.

Golumbic, E. M. Z., N. Ding, S. Bickel, P. Lakatos, C. A. Schevon, G. M. McKhann, R. R. Goodman, R. Emerson, A. D. Mehta and J. Z. Simon (2013). “Mechanisms underlying selective neuronal tracking of attended speech at a “cocktail party”.” Neuron 77(5): 980–991.

Greenlee, J. D., R. Behroozmand, C. R. Larson, A. W. Jackson, F. Chen, D. R. Hansen, H. Oya, H. Kawasaki and M. A. Howard, 3rd (2013). “Sensory-motor interactions for vocal pitch monitoring in non-primary human auditory cortex.” PLoS One 8(4): e60783.

Howell, P. and A. Archer (1984). “Susceptibility to the effects of delayed auditory feedback.” Percept Psychophys 36(3): 296–302.

Mesgarani, N. and E. F. Chang (2012). “Selective cortical representation of attended speaker in multi-talker speech perception.” Nature 485(7397): 233–236.

Niziolek, C. A., S. S. Nagarajan and J. F. Houde (2013). “What does motor efference copy represent? Evidence from speech production.” Journal of Neuroscience 33(41): 16110–16116.

Yates, A. J. (1963). “Delayed auditory feedback.” Psychol Bull 60: 213–232.

